# The ATPase activity of *E. coli* RecA prevents accumulation of toxic complexes formed by erroneous binding to undamaged double strand DNA

**DOI:** 10.1101/361097

**Authors:** Daniil V. Gataulin, Jeffrey N. Carey, Junya Li, Parisha Shah, Jennifer T. Grubb, Douglas K. Bishop

## Abstract

The *E. coli* RecA protein catalyzes the central step of homologous recombination using its homology search and strand exchange activity. RecA is a DNA-dependent ATPase, but its homology search and strand exchange activities are independent of its ATPase activity. ATP hydrolysis converts a high affinity DNA binding form, RecA-ATP, to a low affinity form RecA-ADP, thereby supporting an ATP hydrolysis-dependent dynamic cycle of DNA binding and dissociation. We provide evidence for a novel function of RecA’s dynamic behavior; RecA’s ATPase activity prevents accumulation of toxic complexes caused by direct binding of RecA to undamaged regions of dsDNA. We show that a mutant form of RecA, RecA-K250N, previously shown to be toxic to *E. coli*, is a loss-of-function ATPase-defective mutant. We use a new method for detecting RecA complexes involving nucleoid surface spreading and immunostaining. The method allows detection of damage-induced RecA foci; STED microscopy revealed these to typically be between 50 and 200 nm in length. RecA-K250N, and other toxic variants of RecA, form spontaneous DNA-bound complexes that are independent of replication and of accessory proteins required to load RecA onto tracts of ssDNA *in vivo*, supporting the hypothesis that RecA’s expenditure of ATP serves an error correction function.

## INTRODUCTION

Homologous recombination is a mechanism used by all organisms to repair DNA lesions such as double-stranded breaks (DSBs) and collapsed replication forks (1-3). Inability to repair these lesions results in cell death, genetic instability, birth defects, and cancer (4, 5). To carry out its function in homologous recombination, RecA must assemble into helical nucleoprotein filaments on the ssDNA tracts formed at sites of double strand DNA breaks or stalled replication forks. Although RecA binds rapidly to ssDNA in purified system, its ability to bind tracts of ssDNA *in vivo* requires one of two pathways that allow RecA to outcompete the high affinity ssDNA binding protein SSB (6, 7). One of these pathways requires the multifunctional recombination protein RecBCD (8); the other pathway requires the assembly mediator protein RecFOR (9). Once assembled, RecA-ssDNA filaments search genomic dsDNA for a sequence complementary to that of the bound ssDNA. Identification of a homologous region is coupled to a strand exchange reaction that forms hybrid dsDNA. Hybrid dsDNA tracts template the repair process so that broken chromosomes and stalled forks are healed without insertion or deletion of DNA sequences (1).

RecA is a DNA-dependent ATPase (1, 10, 11). The ATP bound form of RecA has a higher affinity for ssDNA and dsDNA than does the ADP-bound form (12, 13). This difference in affinity, combined with the protein’s DNA-dependent ATPase activity, results in a dynamic DNA binding cycle: RecA-ATP cooperatively binds DNA to form helical nucleoprotein filaments; ATP hydrolysis throughout the filament converts RecA-ATP to RecA-ADP; and RecA-ADP protomers dissociate from DNA. Because protomer-protomer interactions within the filament stabilize DNA binding, dissociation of RecA-ADP protomers occurs predominantly from filament ends (14, 15).

The discovery of the relationship between RecA’s ATPase activity and its DNA binding affinity, together with the demonstration that strand exchange can occur in the absence of ATP hydrolysis, led to the proposal that a function of RecA’s ATPase activity in cells is to release the protein from the products of strand exchange, thereby making the hybrid intermediate available to late-acting recombination proteins (16, 17). RecA’s ATPase activity has been shown to have a variety of functions in biochemical systems including: strand exchange across short heterologous sequences (18); redistribution of RecA monomers within the presynaptic filament (19, 20); limiting filament length (21); and allowing release of RecA-ssDNA filaments that otherwise stall as homology-dependent synaptic complexes at regions of limited homology (22). In addition, ATP hydrolysis is coordinated along RecA filaments allowing filaments to function as rotary motors that drive long-range strand exchange (11, 23).

Although these prior studies have identified various biochemical activities associated with RecA-mediated ATP hydrolysis *in vitro*, relatively little is known about the function(s) of the ATPase activity *in vivo*. However, one important pair of studies sought to address this issue. In the first study, a set of RecA mutants was identified that are toxic to *E. coli* (24). One of these toxic variants, RecA-E96D, was shown to be DNA binding and recombination proficient, but to lack ATPase activity (25). These findings led Campbell and Davis to propose that the ATPase activity of RecA allows release of the protein from recombination intermediates, with failure of release in RecA-E96D expressing cells causing toxicity. However, the authors noted key limitations to the interpretation of their data. First, they noted that it was not possible to definitively determine if the toxicity associated with RecA-E96D resulted from the protein’s defective ATPase activity because the protein displays abnormally fast DNA binding that might cause the observed toxicity. Second, they recognized that their results did not allow them to determine if the toxicity they observed was associated with defective attempts at recombinational repair, or some other mechanism (25).

Our prior work on the eukaryotic homologs of RecA, Rad51 and Dmc1, led us to consider an alternative to Campbell and Davis’s preferred explanation for the toxicity of RecA-E96D; we hypothesized that the toxicity results from direct binding of the protein to undamaged dsDNA regions of the *E. coli* chromosome. We have shown that budding yeast Rad51 and Dmc1, as well as human RAD51, form complexes on undamaged chromatin that can become genotoxic unless removed by a RAD54-family dsDNA-specific, ATP-hydrolysis-dependent, translocase (26-28). RAD54 family translocases use the energy of ATP hydrolysis to displace RAD51 and DMC1 from duplex DNA (29,30). RAD51 and DMC1 display much lower intrinsic DNA-dependent ATPase activity than RecA (*k*_cat_ 0.5-1.0 min^−1^ compared to 30 min^−1^ (31), likely explaining why eukaryotic cells rely on the translocases for strand exchange protein binding dynamics.

In this work, we biochemically characterize another toxic RecA variant, RecAK250N, which was also discovered in the screen for toxic RecA variants (24). We show that, unlike RecA-E96D, RecA-K250N does not display higher than normal DNA binding activity, allowing us to ascribe its toxicity to the ATPase defect. We describe a new method for detecting nucleoid-bound structures of RecA in *E. coli* cells that have suffered DNA damage and use super-resolution light microscopy to measure the lengths of RecA filaments formed following gamma irradiation. We then use the method to detect the DNA-bound structures formed by RecA-E96D and RecA-K250N. We provide evidence that these structures are very unlikely to be recombination or replication intermediates, although their ability to bind nucleoids and confer toxicity depends on DNA binding. These findings provide further support for our hypothesis that the ATP hydrolysis-dependent DNA binding dynamics of RecA-family strand exchange proteins prevents their toxic accumulation on undamaged chromosomes.

## MATERIALS AND METHODS

### Bacterial strains, plasmids, and growth conditions

*E. coli* strains used in this study are listed in Supplementary Table 1. DNA oligos used for site-directed mutagenesis and construction of 6xHis-tagged strains are listed in Supplementary Table 2.

Unless otherwise specified, cultures were grown with shaking for 12-16 hours (overnight) at 37°C, then diluted 50-100-fold to OD_600_ = 0.05 and incubated at 37°C. Experiments were conducted when cultures were in log phase having reach a density of OD_600_ = 0.2-0.4.

### Growth media, DNA substrates, chemicals

TB and LB medium were previously described (24). Minimal aspartate and K media were also previously described (32). For arabinose induction experiments media were supplemented with L-(+)-Arabinose (Sigma) to 0.2%. Where appropriate, antibiotics were used at the following concentrations: Kanamycin, 50 μg/mL; Ampicillin, 100 μg/mL; Tetracycline, 10 μg/mL; Chloramphenicol, 17 μg/mL. ATP was purchased from Sigma. ATPγS was purchased from Cayman Chemical.

5’-Fluorescein-labeled RG-1(+) ssDNA (84 nt) used in Fluorescent polarization assays is described in (33) and was purchased from Eurofins. Preparation of fluorescein-labeled 162 bp dsDNA used has been described (34). This duplex substrate contains covalently-closed hairpin ends that prevent transient opening of ends that might allow RecA to bind the resulting ssDNA tracts.

### Purification of RecA proteins

Overexpression of *recA* variants was carried out as previously described (24), except 4 L of culture was used instead of 9 L. After induction, cells were harvested by centrifugation, washed with Buffer P (25 mM sodium phosphate, pH 7.5; 10% glycerol, 1 M NaCl, 50 mM imidazole,1 mM DTT), pelleted by centrifugation and stored at −80°C. Non-toxic *recA* alleles were cloned into pET-9c vector and expressed in BLR(DE3) pLysS cells. Several colonies were inoculated into 100 mL of LB supplemented with 1% glucose and 0.5% NaCl and allowed to incubate at room temperature without shaking overnight (12-14 h). Cultures were then transferred to 37°C shaking incubator, grown to OD_600_ = 0.6, back-diluted into 1 L of LB (1% glucose, 0.5% NaCl) to OD_600_ = 0.05, and grown at 37°C to OD_600_ = 0.8, at which point IPTG was added to 0.5 mM to induce RecA expression. After 1 h of induction, temperature was lowered to 30°C and cultures were incubated for another hour. After induction, cells were harvested by centrifugation, washed with Buffer P, and the pellet was stored at −80°C.

Cell lysate was prepared by resuspending pellet in 40 mL cell buffer P containing EDTA-free protease inhibitor cocktail (Roche) using Dounce homogenizer and passed through French Pressure Cell Press (American Instrument Company) three times at 18,000-20,000 psi. The lysate was centrifuged at 40,000 rpm (Beckman Ti70 rotor) and clear supernatant was saved. The supernatant was added to 1 mL of Ni Sepharose resin (GE Healthcare) that was pre-equilibrated with Buffer T and incubated at 4°C for 1 h to allow protein to bind to the resin. The resin was pelleted by centrifugation, washed twice with buffer P, and resuspended in 5 mL of buffer T (25 mM Tris-HCl, pH 7.5, 10% Glycerol, 500 mM NaCl, 1 mM DTT) with 30 mM imidazole. The mixture was loaded into a protein elution column (1.5 cm diameter), washed with 40 mL of Buffer T with 30 mM imidazole, followed by another wash with 20 mL of Buffer T with 50 mM imidazole. The protein was eluted in 1 mL fractions with 5 mL of Buffer T with 80 mM imidazole, 10 mL of Buffer T with 300 mM imidazole, and 5 mL Buffer T with 500 mM imidazole. OD280 was measured for each fraction using NanoDrop ND-1000 spectrophotometer and peak fractions were pooled. Purified protein was exchanged into a Storage Buffer (25 mM Tris-HCl, pH 7.5, 0.1 mM EDTA, 1mM DTT, 50 mM NaCl, 20% glycerol) using PD-10 desalting columns (Amersham Biosciences) and stored at −80°C. Purity of the protein was assayed by SDS PAGE and concentration was determined using absorbance at 280 nm and extinction coefficient of 22,300 M^−1^ cm^−1^ (35).

### ATPase assays

RecA ATPase activity was measured using Malachite Green assay (36). Reactions contained 25 mM Tris-HCl (pH 7.5), 1 mM ATP, 4 mM MgCl_2_, 1 mM DTT, 100 μg/mL BSA, 1 μM RecA, and 36 nM pBS90 ssDNA or 16.7 nM pRS306 dsDNA. After incubation for indicated times at 37°C, dye reagent (0.02% Malachite Green, 0.3% Na_2_MoO_4_, 0.8N H_2_SO_4_, 0.04% Tween) was added and the mixture was incubated for 5 min, after which time A_635_ was measured using Tecan Infinite F200 Pro plate reader. The data was plotted as A_635_ vs time.

### Fluorescence Polarization Experiments

Fluorescence polarization (FP) was used as described in (34) to measure RecA binding affinity and kinetics of interaction with ssDNA and dsDNA. Reactions contained 25 mM Tris-HCl (pH 7.5), 3 mM ATP, 10 mM (ssDNA experiments) or 1 mM (dsDNA experiments) MgCl_2_, 1 mM DTT, 100 μg/mL BSA, ATP regeneration system (8 mM phosphocreatine, 10 U/mL creatine phosphokinase). Reactions were initiated by the addition of RecA (diluted to appropriate concentration in RecA storage buffer) and FP measurements were taken on Tecan Infinite F200 Pro plate reader at 37°C. For DNA binding experiments, reactions were pre-incubated for 20 min at 37°C before data collection. For RecA on-rate experiments, reaction components were chilled on ice and FP data collection was started immediately after addition of RecA protein. FP measurements were taken at room temperature every 10 sec for 30 min (ssDNA) or 20 sec for 45 min (dsDNA). For RecA dissociation-rate experiments, RecA filaments were assembled on fluorescein-labeled ssDNA or dsDNA at 30°C for 30 min and FP data collection was started immediately after addition of 10-fold excess competitor ssDNA. FP measurements were taken at 30°C every 10 sec for 30 min (ssDNA) or 20 sec for 45 min (dsDNA). Competitor DNA used was a 90-mer with the following sequence: 5’-GGCACCAACA CAAAACACAT CTACACTCAA CAATCACTTT TTATACAACA CTTCTTCTCT CACATACAAC ACTTCTGGCA CCAACACAAA.

### D-loop Assay

A modified displacement loop (D-loop) assay used here replaces ^32^P-ssDNA with fluorescently labeled ssDNA substrate (37). D-loop formation was achieved by incubating 1.5 μM RecA protein in a 10 μL buffer containing 70 mM Tris-HCl (pH 7.5), 10 mM MgCl_2_, 5 mM DTT, 100 μg/mL BSA, 3 mM ATP, ATP regeneration system (10 mM phosphocreatine, 10 U/mL creatine phosphokinase), 30nM 5’Quasar670-modified 90-mer ssDNA oligo (5’AAAAGGCCTC TAGGTTCCTT TGTTACTTCT TCTGCCGCCT GCTTCAAACC GCTAACAATA CCTGGGCCCA CCACACCGTG TGCATTCGTA, purchased from LGC BioSearch Technologies), and 5 nM homologous pRS306 dsDNA for 15 min at 37°C. The rest of the protocol was carried exactly as described in (37). Quantity One (BioRad) software was used to analyze gel images, and D-loop yield was expressed as a percentage of input plasmid DNA.

### Electron Microscopy

RecA filament formation was assayed by incubating 4 μM RecA in a buffer containing 25 mM Tris-HCl (pH 7.5), 10 mM MgCl_2_, ATP regeneration system (8 mM phosphocreatine, 10 U/mL creatine phosphokinase), 1 mM DTT, 3 mM ATP, 3% (v/v) glycerol, 7.5 mM NaCl, and 60 nM no-fold 90C ssDNA for 10 min at 37°C. Filaments were stabilized by further addition of 3 mM ATPγS and 3 min incubation at 37°C. The samples were spotted onto carbon coated grids and negatively stained with 1% uranyl acetate. Images were collected using GATAN digital camera on FEI Tecnai F30 scanning transmission electron microscope at 49,000X magnification.

### Gamma and UV Irradiation

Logarithmically growing cultures (OD_600_ = 0.2-0.4) were transferred to 50 mL conical tubes and chilled on ice. Chilled cultures were irradiated with 280 Gy gamma rays from a GammaCell ^60^Co source (Atomic Energy of Canada Ltd., Kanata, Ontario), transferred to clean flasks, and returned to growth conditions. After 30 min of incubation, samples were taken for chromosome spreading. For UV irradiation experiments, 10 mL of logarithmically growing cultures were transferred to 100 mm plastic petri dish and irradiated to 20 J/m^2^ on a rotating platform. Cultures were returned to growth conditions for 60 min prior to harvesting for chromosome spreading.

### Induction of RecA variants

Cells harboring different *recA* alleles under the control of arabinose-inducible promoter were grown to early log phase in K medium, pelleted, and resuspended in glucose-free K medium with 0.2% arabinose for induction of RecA protein. After 30 min induction under normal growth conditions, cells were harvested for chromosome spreading and Western blotting. *dnaAts* cultures were grown at 30°C and were pre-incubated at 42°C for 90 min prior to induction.

### Western blotting

Cells from 1 mL culture were pelleted, resuspended in 100 μL SDS lysis buffer (50 mM Tris-HCl pH 6.8, 2% SDS, 10% glycerol, 0.1% bromophenol blue, 5% 2-mercaptoethanol, 50 mM dithiothreitol), and boiled for 5 min. After boiling, suspension was pelleted and 15 μL of supernatant was run on a 10% acrylamide (37.5:1 acrylamide:bis) SDS-PAGE gel by standard methods. Protein was transferred to Immobilon-P membrane (Millipore) using a BioRad Mini-PROTEAN 3 Trans-Blot electrophoretic transfer cell. Blots were incubated at 4°C for 12-14 h with monoclonal mouse anti-RecA antibody diluted to 0.11 ng/mL in blocking buffer. Secondary labeling was done using horseradish peroxidase conjugated sheep anti-mouse antibody (GE Healthcare) diluted 1:1,000 in blocking buffer. Blots were treated with Western Lightning Plus chemiluminescence reagent (Perkin Elmer) and imaged using the Alpha Innotech FluorChem SP imaging system.

### Nucleoid spreading

5 mL cultures were harvested and pelleted. The supernatant was discarded and the pellet kept on ice until ready for processing. Pellets were resuspended in 0.45 mL of osmotic shock incubation buffer (10 mM NaH_2_PO_4_, 10 mM EDTA, 100 mM NaCl, and 30% sucrose) (38). 50μL of a freshly made 4 mg/mL lysozyme solution in osmotic shock incubation buffer was added to each sample. Samples were briefly and gently mixed and incubated at room temperature for 30 min to generate spheroplasts. After incubation, 10 μL of spheroplast suspension was deposited on the center of a glass cover slip (24#x00D7;50−1.5, Fisher), which was secured to a microscope slide with cellophane tape, and 40 μL of paraformaldehyde solution (3% paraformaldehyde, 3.4% sucrose, pH 7.0) was added to the spheroplast suspension. The liquid was distributed on the cover slip surface with a glass rod. Spheroplasts were lysed by adding 1 mL of 10 mM sodium chloride to the liquid on the cover slip and an additional 80 μL of paraformaldehyde solution was added to the lysate. Cover slips were allowed to air dry at room temperature overnight. If not used immediately, cover slips were stored at −20°C.

### Immunostaining

Cover slips were immersed in 0.2% Photo-Flo 200 (Kodak) for 30 sec then transferred to 1x TBS (140 mM sodium chloride, 2.7 mM potassium chloride, 25 mM Tris-Cl pH 8.0) for 5 min. Excess liquid was drained from the cover slips and 350 μL blocking buffer (1x TBS plus 5% non-fat dry milk (Bio-Rad) and 0.1% TWEEN 20) was added to the cover slip surface. Cover slips were incubated in a damp chamber for 15 min at room temperature. Excess blocking buffer was drained from the cover slips and 40 μL primary antibody diluted in blocking buffer was added and covered with parafilm strip to prevent drying out. Monoclonal mouse anti-RecA (MBL) was used at 0.11 ng/mL. Rabbit anti-SSB (gift from Ute Curth at Medizinische Hochschule Hannover) was used at a 1:20,000 dilution. Cover slips were incubated with primary antibody in a wet chamber for 12-16 h at 4°C. Parafilm was removed and the cover slips were washed twice in 1x TBS for 10 min each. Excess liquid was drained from the surface and 40 μL secondary antibody diluted in blocking buffer was applied as before. Secondary antibodies were Alexa Fluor 488 goat anti-mouse IgG (Invitrogen) for RecA and Alexa Fluor 594 goat anti-rabbit IgG (Invitrogen) for SSB, both used at 2 ng/mL. Cover slips were incubated with secondary antibody in a wet chamber for 2 h at 4°C and washed as before. After secondary staining, cover slips were dried at room temperature and then mounted onto microscope slides with Vectashield mounting medium with DAPI (Vector Laboratories H-1200) and sealed with clear nail protector. Mounted slides were stored at 4°C.

### Microscopy and image analysis

Epifluorescence microscopy was conducted using a Zeiss Axio Imager.M1 fluorescence microscope as previously described (39). STED microscopy was conducted using a Leica TCS STED CW instrument. Samples for STED microscopy were prepared and stained as for epifluorescence microscopy except that STED samples were prepared directly on 22#x00D7;22−1.5 cover slips and were mounted using ProLong Gold (Invitrogen). Image processing and analysis was conducted using NIH ImageJ. Foci were scored manually or using custom-written ImageJ macro (available for download at https://github.com/bbudke/BB_macros/tree/master/Cytology_modules) (no significant difference in focus counts was observed when counted manually or using macro). Statistical comparison of focus data was carried out using a two-tailed Mann-Whitney test for non-normal distributions with significance established at P < 0.05.

### Toxicity assay for mutant *recA* alleles

Toxicity assays of different *recA* mutants were carried as described (24). Briefly, overnight cultures were diluted 100-fold into TB media containing 0.02% L-arabinose and incubated for 3 h at 37°C. Control cultures were diluted into arabinose-free media. Cell numbers were normalized and 5 μL of serial 10-fold dilutions were spotted on LB plates containing appropriate antibiotic. Plates were photographed the next day.

## RESULTS

### Identification of an ATPase defective allele of RecA that does not display abnormally high DNA binding

In order to determine if blocking RecA’s ATPase activity causes the protein to become genotoxic, it is important to identify an ATPase-defective version of the protein that does not display abnormally high DNA binding. The most extensively studied toxic variant of RecA, RecA-E96D, was previously reported to display faster than normal binding kinetics and is therefore unsuitable for use in making genetic inferences regarding the normal *in vivo* function of RecA’s ATPase activity (25). The majority of RecA ATPase deficient mutants identified previously were unsuitable for the study because they are DNA binding-deficient (40). However, previous studies indicated RecA-K250N and RecA-K250R are DNA binding proficient, ATPase defective variants (41, 42). Expression of RecA-K250N was reported toxic, like RecA-E96D (24), although it has not previously been tested for its rate of binding nor its ability to bind dsDNA. RecA-K250R was reported to reduce cell growth rate and to be partially defective in ATPase activity (41).

We tested ATPase activity in the presence of either a 84-nt oligonucleotide (ssDNA) or a 162-nt duplex (dsDNA) for four variants of RecA with a Malachite Green assay that reported *P_i_* release (36, 43). Wild type RecA hydrolyzed ATP with the greatest apparent rate reaching completion in about 30min in the presence of either ssDNA or dsDNA. The K250R mutant hydrolyzed ATP with an intermediate rate while E69D and K250N displayed almost no ATPase activity in either ssDNA or dsDNA, consistent with previous observations (25, 41) (Figure 1A, 1B). As expected, almost no ATPase activity was observed for any of the RecA variants in the absence of DNA (K250N and to a lesser extent K250R showed low levels of ATPase activity at 60 minutes) (Supplementary Figure 1).

**Figure 1.**
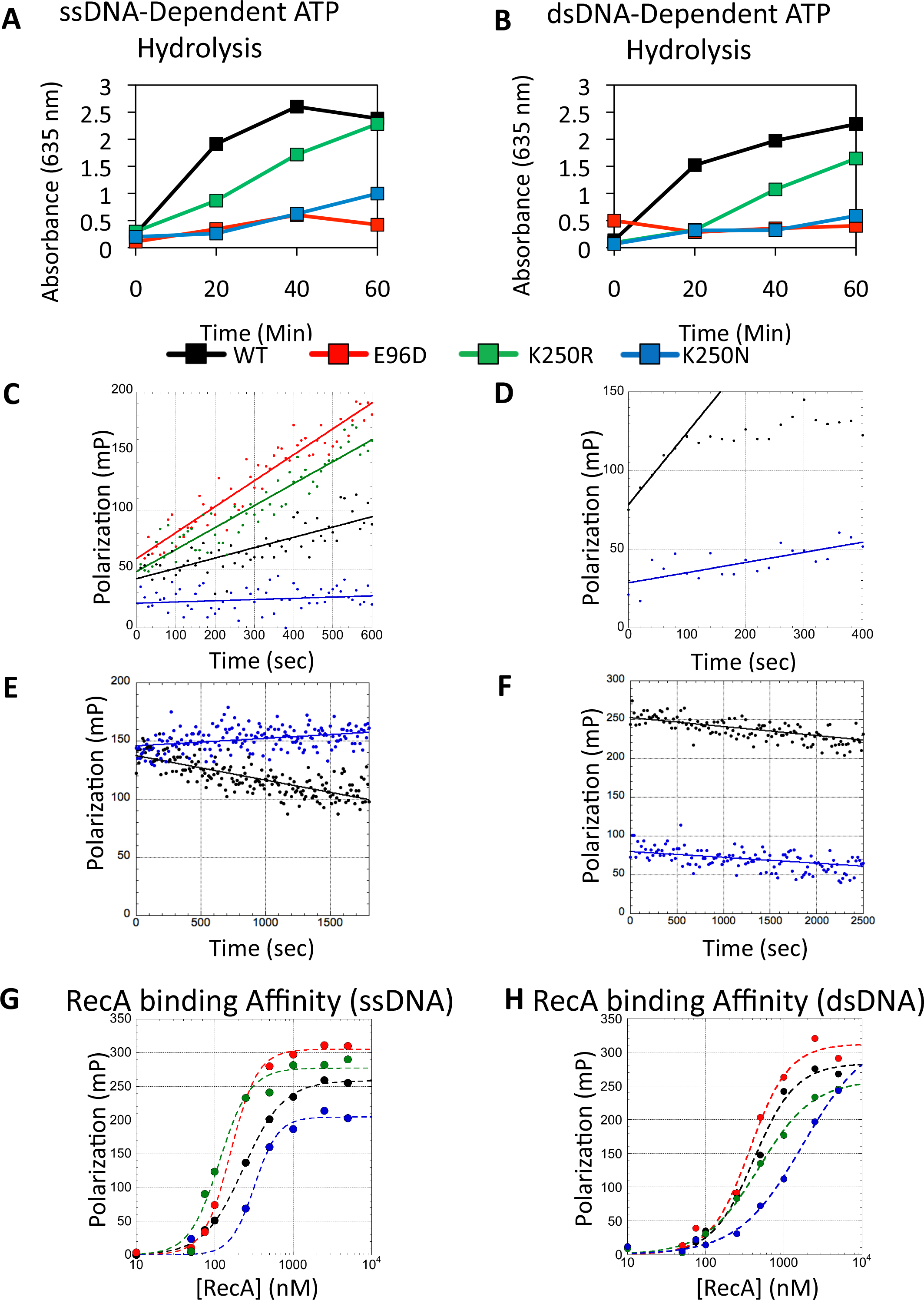
Biochemical characterization of RecA ATPase-defective variants. A. Malachite green ATPase assays with 36 nM 90-mer ssDNA oligonucleotide. B. Same as A. except DNA was 16.7 nM 200 bp pRS306 dsDNA (position 654-853). C. Kinetic analysis of DNA binding measured by fluorescence polarization. DNA substrate was 2.4 nM RG-1(+) ssDNA. RecA was used at 250 nM. Reaction components were chilled on ice and FP data collection was started immediately after addition of RecA. D. Same as C except DNA substrate was 4.8 nM 162 bp dsDNA with hairpin ends (see Methods and Materials) and RecA was used at 2.5 µM. E. RecA turnover from ssDNA. RecA (0.7 µM) was pre-assembled on 2.4 nM RG-1(+) ssDNA at 30°C and data collection was started immediately after addition of 22 nM competitor ssDNA. F. Same as E except DNA substrate was 4.8 nM 162 bp “dumbbell” dsDNA, RecA was used at 1 µM. G. Equilibrium analysis of DNA binding via fluorescence polarization. Reactions we pre-incubated at 37°C for 30 min. DNA substrate was 2.4 nM RG-1(+) ssDNA. H. same as E except DNA was 4.8 nM 162 bp hairpin dsDNA.

We next measured the rate of association between RecA variants and either ssDNA or dsDNA by employing a fluorescence polarization assay. The wild type enzyme associated with the ssDNA oligonucleotide with an initial rate of 0.07 s^−1^ whereas the RecA-E96D and RecA-K250R variants displayed association rates of 0.22 and 0.17 s^−1^, respectively, with ssDNA. The RecA-K250N variant showed limited ssDNA association with an initial rate of 0.01 s^−1^ (Figure 1C). We next compared the association rates of RecA-WT to RecA-K250N on dsDNA and found that RecA-WT displayed an initial rate of association of 0.46 s^−1^ whereas the RecA-K250N variant associated with dsDNA at a significantly slower with a rate of 0.07 s^−1^ (Figure 1D). We also measured dissociation rates of RecA-WT and RecA-K250N from both ssDNA and dsDNA. Unlike RecA-WT, which dissociated from ssDNA at an average rate of 0.021 s^−1^ upon addition of competitor DNA, no RecA-K250N dissociation was observed under the same conditions (Figure 1E). When dsDNA was used as a substrate, rate of RecA-WT dissociation was 0.012 s^−1^ whereas RecA-K250N dissociated with an average rate of 0.0075 s^−1^ (Figure 1F).

Additionally, we tested the RecA-DNA binding under equilibrium conditions to determine affinity constants (*K_d_*) for each RecA variant with either ssDNA or dsDNA. The E96D and K250R mutants bound ssDNA with the highest affinity (*K_d_* = 150 ± 10 and 110 ± 10 nM, respectively) consistent with the observations that these variants bound ssDNA more quickly and hydrolyzed ATP more slowly than wild type RecA. The K250N variant bound ssDNA with lower affinity (*K_d_* = 300 ± 40 nM), about 1.5-fold weaker than wild type (*K_d_* = 210 ± 20 nM) consistent with the observations that the K250N mutation significantly impaired DNA association (Figure 1G). Wild type, E96D and K250R mutants all bound dsDNA with similar affinities (*K_d_* = 340-470 nM), but the K250N mutant bound with a significantly lower (*K_d_* > 1000 nM), but easily detectable affinity (Figure 1H).

As a further control against gross structural or functional defects that are independent of observed ATPase defects, we tested the ability of each mutant protein to carry out homology search and strand exchange using a D-loop assay. D-loop assay employs a ssDNA oligonucleotide and a supercoiled plasmid containing a sequence corresponding to the oligonucleotide; here we used fluorescently-labelled ssDNA rather than the more traditional radiolabeled substrate (37). These experiments revealed that versions of RecA carrying the E96D, K250N, or K250R mutations all displayed substantial levels of D-loop activity, with RecA-E96D and RecA-K250R displaying higher levels of D-loop activity than RecA-WT and RecA-K250N displaying lower levels than WT, consistent with binding data (Supplementary Figure 2A). Although it has no bearing on the conclusions of the paper, we should note that our preparation of RecAE96D appears to have a minor topoisomerase contamination. We also examined the structure of filaments by electron microscopy of negatively stained samples and found all mutants form filaments on ssDNA that have the expected diameter and helical pitch (Supplementary Figure 2B). These results indicate that all the variants examined are properly folded and active.

In summary, our biochemical analyses show that RecA-K250N meets the criteria for testing the normal *in vivo* function of RecA’s ATPase activity via loss-of-function genetic analysis; RecA-K250N is an ATP hydrolysis-defective mutant that retains DNA binding and D-loop activity, but does not bind DNA more rapidly than RecA-WT. This result implies that loss of ATPase activity is sufficient to result in RecA-mediated toxicity; enhanced DNA binding is not required.

### Bacterial chromosome spreading allows detection of damage-induced RecA and SSB foci

Immunostaining of spread chromosomes from eukaryotic cells has proven a convenient and powerful method for analyzing the chromosome binding behavior of proteins involved in homologous recombination (44). To develop a similar approach to study proteins bound to *E. coli* chromosomes, we combined the *E. coli* nucleoid isolation technique of Woldringh and colleagues (38) with the chromosome spreading method of Klein and Loidl (45) that was originally developed to visualize meiotic budding yeast chromosomes (see Methods for details). To evaluate if chromosome-bound RecA complexes could be observed following induction of DNA damage, cells growing logarithmically in rich medium (LB) were exposed to 280 Gy of ionizing radiation from a ^60^Co gamma source. Nucleoid spreads were prepared 30 min after irradiation and co-immunostained with antibodies against RecA and SSB. Following irradiation, a punctate staining pattern was detected for RecA that was not seen in unirradiated controls (Figure 2B). About 47% of spread chromosomes contained at least one RecA focus, and the average number of RecA foci was 2.2 ± 0.4. In contrast, only 2% of spread chromosomes from unirradiated controls contained any obvious RecA staining, and this small subpopulation typically displayed a single focus per nucleoid (Figure 2A). We obtained similar results following UV irradiation: 61% of wild type cells showed RecA focus formation after irradiation with 20 J/m^2^ UV, with an average of 3.3 ± 0.4 RecA foci per focus-positive nucleoid (Figure 2C, 2D). Small numbers of RecA foci in *recA* cells represent non-specific background staining.

**Figure 2.**
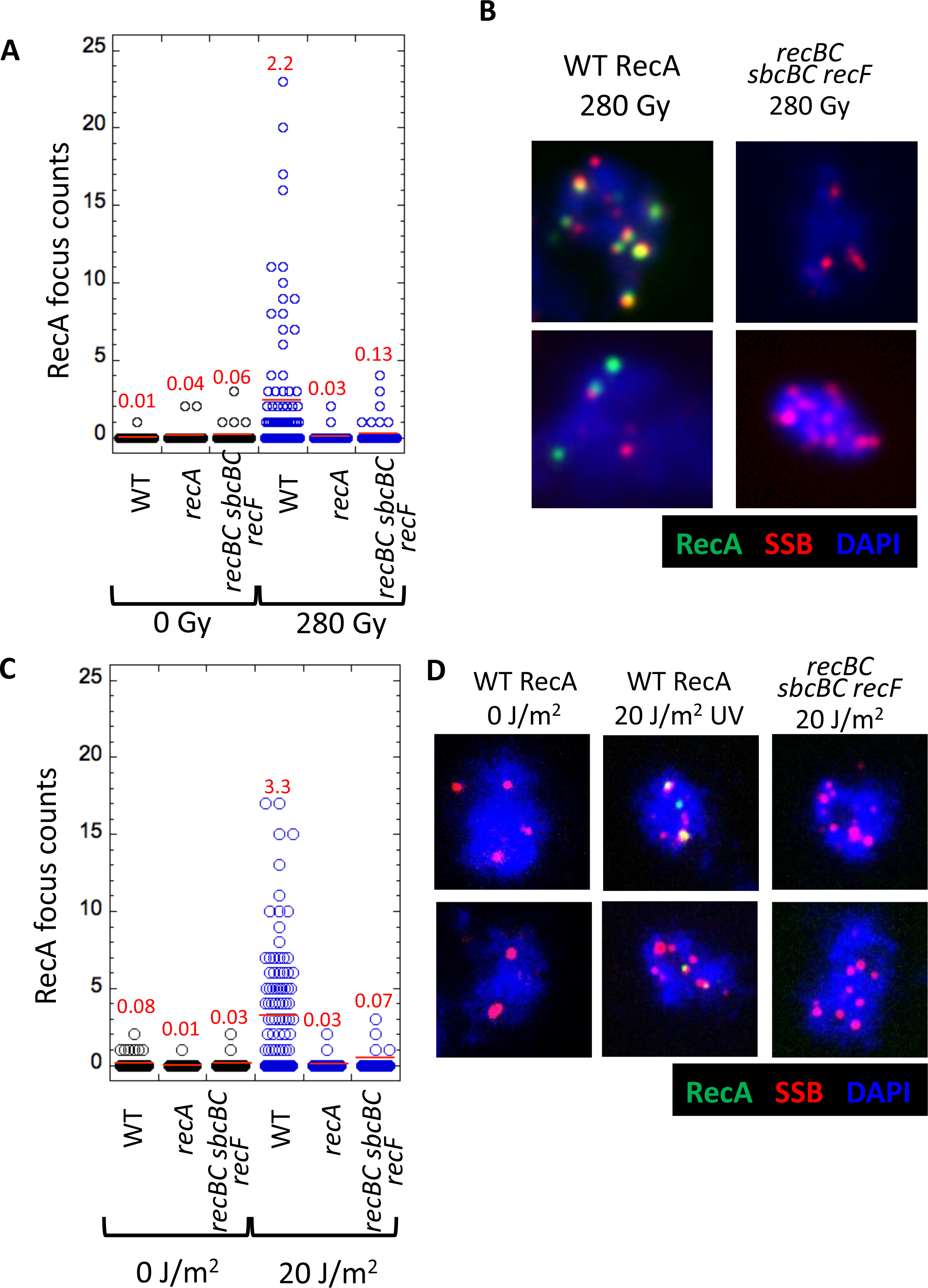
Analysis of damage-induced RecA foci. A. RecA focus counts from spread nucleoids. Log phase cells in LB medium were treated with 280 Gy (blue) and incubated at 37°C for 30 min and then subjected to the nucleoid spreading protocol and indirectly immunostained for RecA and SSB. Control cells (black) were not subject to irradiation. The *recA* null mutant strain is BUG74. N = 100 nucleoids per sample. Average focus counts are shown as red lines and indicated in red font. B. representative nuclei from the experiment shown in A. RecA (green), SSB (red); DNA was stained with DAPI (blue) C. Same as A except cells were treated with 20 J/m^2^ 302 nm UV light. D. representative nuclei from the experiment shown in C.

We next asked if the damage-induced RecA foci we observe depend on functional RecBCD or RecFOR accessory factor complexes. We found that, unlike cells expressing RecBCD and RecFOR, only 7% of spread chromosomes from *recBC sbcBC recF* cells formed RecA foci after 280 Gy of ionizing radiation, with an average of 0.13 ± 0.06 foci per nucleoid; overall the mutant strain displayed 17-fold fewer foci than wild type (Figure 2A). Similar results were obtained by UV (Figure 2B). After treatment with 20 J/m^2^, only 4% of spread chromosomes from *recBC sbcBC recF* cells formed RecA foci, with an average of 0.07 ± 0.04 foci per nucleoid. These results indicate that at least 95% of both gamma ray-induced and UV-induced foci, require at least one of the two biochemically and genetically-defined RecA assembly proteins.

### Super-resolution imaging of RecA foci reveals features of *in vivo* filament structure

To characterize damage-induced RecA foci at higher resolution than afforded by standard wide-field microscopy, we employed the super resolution light microscopy method referred to as stimulated emission depletion (STED) microscopy (46). Using this technique, we were able to achieve a resolution of about 50 nm (defined as full width at half maximum of the smallest RecA structure observed). The images obtained by STED demonstrate more clearly that the colocalizing RecA and SSB foci seen via widefield represent closely adjacent, rather than molecularly colocalized, protein structures (Figure 3A). RecA structures that appeared as diffraction-limited foci by wide-field microscopy were resolved into one of four distinct categories by STED (Figure 3B, n=100 foci): 49% resolved into smaller disks or ovoids, 24% into structures with near uniform linear or curvilinear character, and 13% into small paired puncta. The remaining 14% showed no significant change compared to the confocal image. As the persistence length of a RecA-DNA filament is 450 nm (47), it is possible that curvilinear structures such as those observed do not represent continuous nucleoprotein filaments but rather two or more unresolvable, shorter filaments. Closely positioned filaments or paired puncta could represent RecA structures forming on both sides of a DSB or two small complexes assembled on the same tract of ssDNA (48). The diameters, or contour lengths, of the majority of RecA structures observed by STED were in the range of 50-200 nm. Under the assumption that foci reflect canonical RecA nucleoprotein filaments, which are roughly 2nt/nm, the results suggest that readily detected RecA foci reflect nucleoprotein filaments spanning about 100 to 400 nts (or bps).

**Figure 3.**
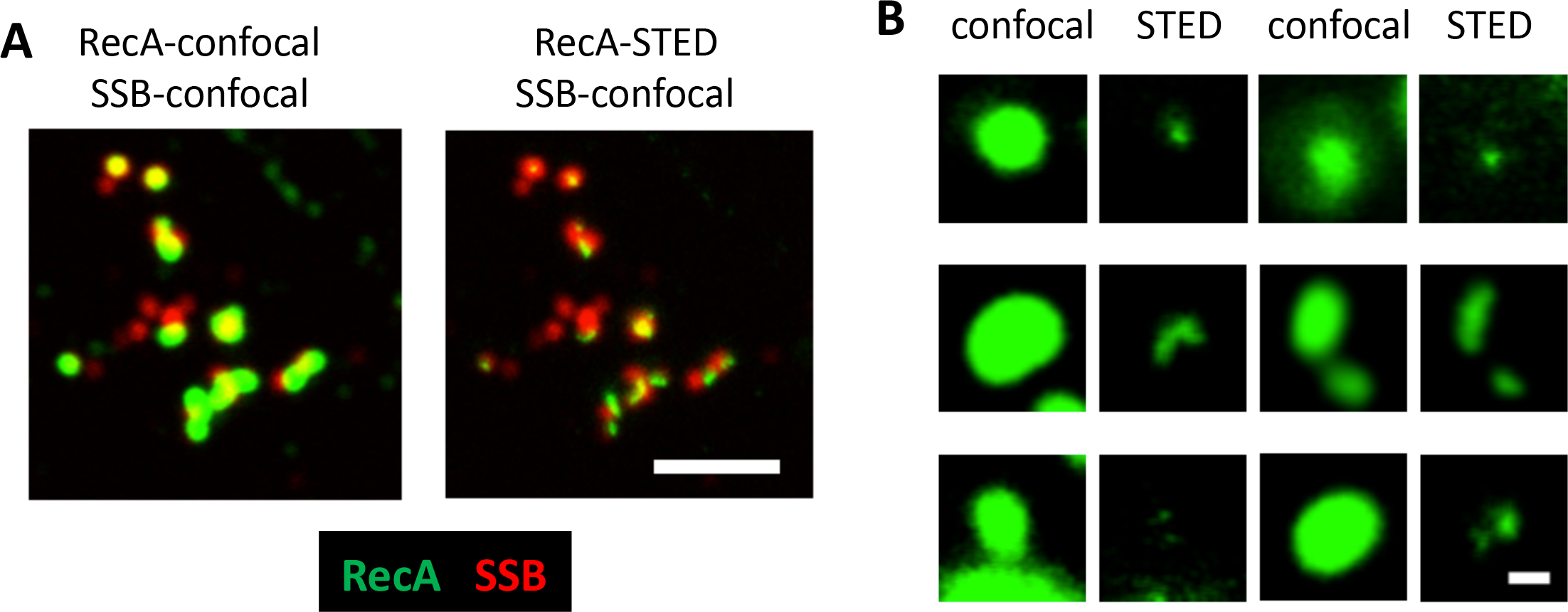
STED imaging of RecA foci. A. Spread nucleoid from a culture (AB1157) treated with 280 Gy gamma rays. Staining was for RecA (green) and SSB (red). Only RecA was imaged in super-resolution, as technical limitations of the STED instrument did not permit super-resolution imaging of both fluors. Note offset localization of RecA and SSB. Scale bar, 2 μm. B. Representative foci viewed by confocal and STED microscopy. Structures appearing as diffraction-limited spots by confocal microscopy often resolved into much smaller puncta (top), linear structures (middle), or pairs of puncta (bottom) when viewed by STED microscopy. Scale bar, 200 nm.

### Toxic RecA mutants form spontaneous complexes on undamaged chromosomes

Having characterized the behavior of wild type RecA protein (RecA-WT) in response to induced DNA damage, we examined the effect of inducing expression of ATPase defective mutant proteins in the absence of any DNA damaging treatment. Specifically, we wanted to ask if RecA’s hydrolytic cycle serves to prevent the protein from accumulating as cytologically-detectable structures on undamaged chromosomes. The expression of each protein was induced from a plasmid in which the coding region of RecA was fused to the arabinose-inducible promoter, P*_araBAD_*. We used Western blotting to determine if the level of expression in this artificially-regulated system was similar to that in cells expressing RecA from the chromosome. We found that the induced level of expression was only about 1.5-fold higher than the level of expression seen in wild type cells, and not as high as the level observed following UV treatment of WT (Supplementary Figure 3). To ensure that the conditions of induction were sufficient to induce toxicity, we tested the cells ability to survive by plating serial dilutions following induction of the protein variants in media containing arabinose. Induction of RecA-E96D and RecA-K250N drastically reduced survival compared to expression of the RecA-WT control, as expected (24). We also found that RecA-K250R did not show a reproducibly-detectable reduction in survival compared to RecA-WT (Figure 4A, 4B).

**Figure 4.**
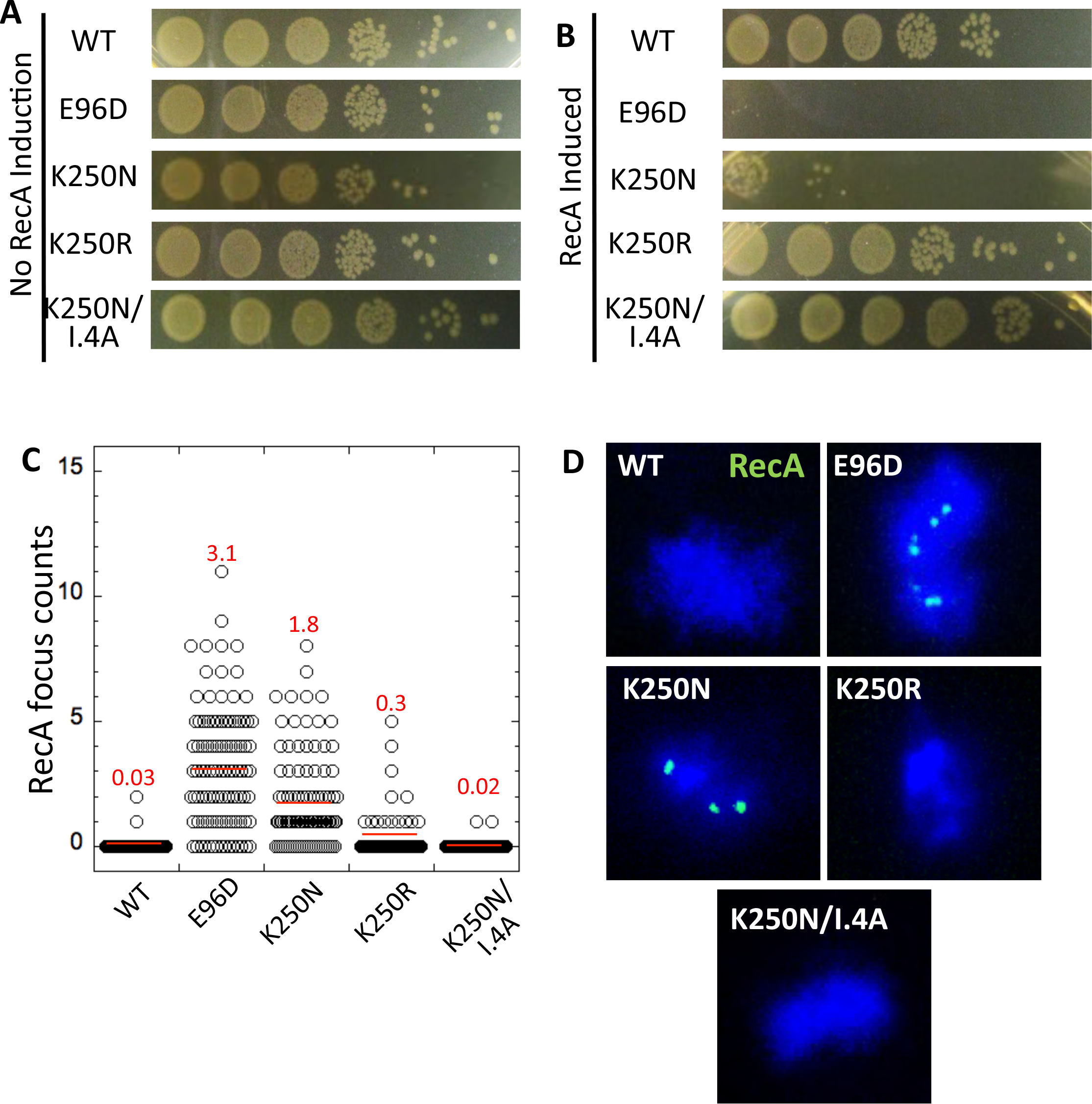
*In vivo* analysis of RecA ATPase-defective variants in strain BNN145. A. Dilution series (10-fold) of aliquots of log phase cultures growing in TB media lacking arabinose. These cultures serve as a control to show similar densities of live cells in each culture. B. Same as A, except cultures were grown for 3 hrs in TB media containing 0.2% L-arabinose to induce RecA expression prior to plating. C. RecA focus counts of cells induced to express RecA variants as in B. n= 100 nucleoids per sample. Average focus counts are shown as red lines and indicated in red font. D. Representative spread nucleoids stained for RecA (green). DNA was stained with DAPI (blue).

Chromosome spreads were then prepared following induction of expression for 30 min and indirectly immunostained for RecA. We found that chromosome spreads from cells expressing RecA-K250N or RecA-E96D displayed spontaneous RecA foci; about 70% of RecA-K250N expressing nucleoids displayed at least one RecA focus with an average of 1.8 ± 0.2 foci/nucleoid; RecA-E96D displayed 3.1 ± 0.24 foci/nucleoid (Figure 4C). In contrast, the frequency of spontaneous RecA-WT foci was much lower, 0.03 foci/nucleoid overall. The frequency of foci seen following induction of RecA-K250R was intermediate between RecA-WT and the other RecA mutant proteins; about 10% of nucleoids contained at least one focus and an average of 0.3 ± 0.1 foci/nucleoid (Figure 4C).

As a control to provide evidence that spontaneous foci represented a DNA-bound species, we designed a mutant variant with 4 alanine substitution mutations within the high affinity DNA binding site of RecA (Site I) in a region of the site referred to as loop 2 (42). We refer to the mutant version of RecA carrying this set of 4 alanine substitutions as RecA-I-4A for DNA binding site I, 4 alanines. Biochemical analysis showed that RecA-I-4A is profoundly deficient in binding both ssDNA and dsDNA in our fluorescence polarization assay (Supplementary Figure 4). In addition, a version of the protein that combined the K250N mutation with the I.-4A mutations also failed to bind DNA *in vitro*. Importantly, only 2% of the RecA K250N/I-4A cells contained spontaneous foci, with overall average of 0.02 foci/nucleoid. Furthermore the I.-4A mutations fully rescued the cellular toxicity associated with K250N (Figure 4A, B). A RecA-E96D/1I-4A mutant gave equivalent results to RecA -K250N/I-4A (Figure 3, Supplementary Figure 5). These results have two implications; first, they suggest that the spontaneous foci formed by RecA-K250N and RecA-E96D represent DNA bound structures, and second, they provide evidence that the toxicity conferred by these proteins is a consequence of DNA binding.

**Figure 5.**
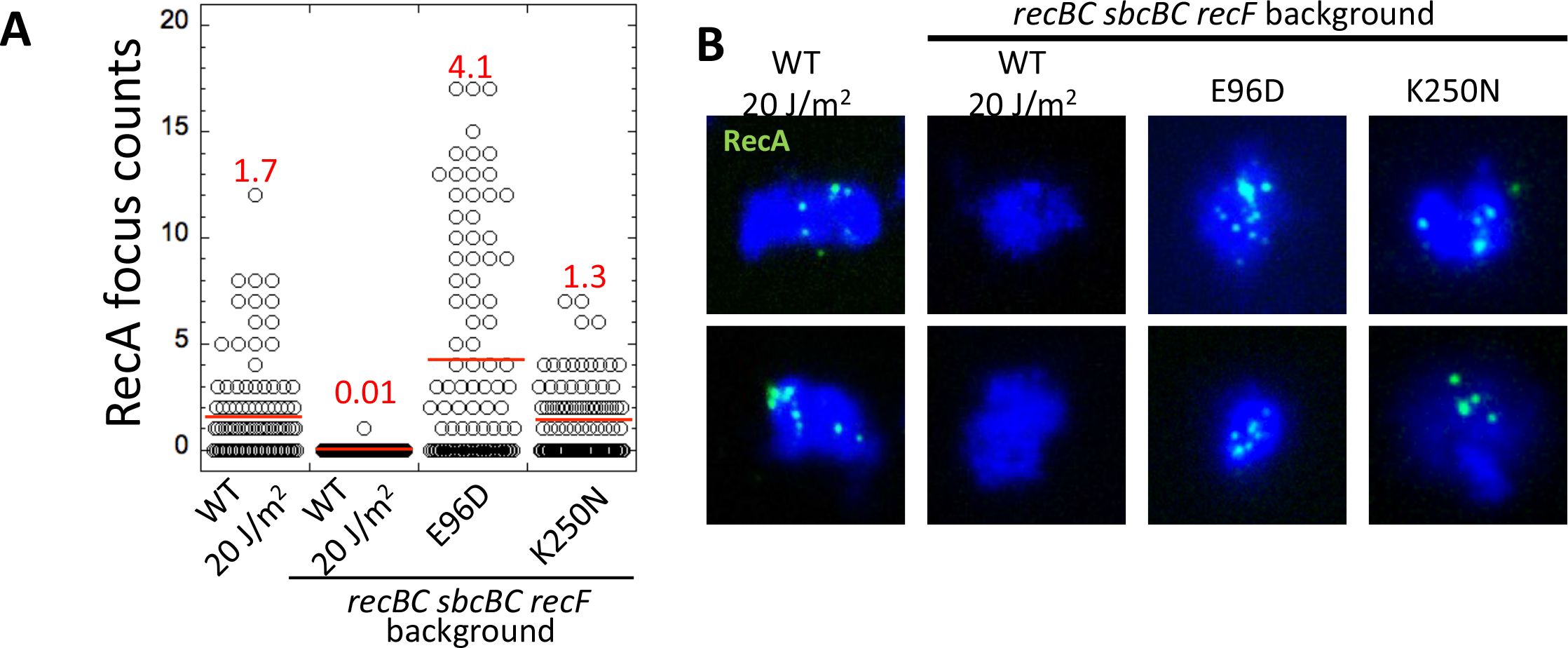
Analysis of RecA ATPase defective variants in *recBC recF* cells (STL2424). A. Spread nuclei were prepared after arabinose induction, foci from individual nucleoids were counted, and data plotted as described in legend of Fig. 4. B. Representative nucleoids from the experiment shown in A. RecA (green); DNA was stained with DAPI (blue).

Having shown that the toxic forms of RecA form spontaneous foci, we wished to determine if these foci represent ssDNA-bound forms (as expected if they represent recombination intermediates) or if they result from binding to undamaged dsDNA regions of the chromosome. To address this question, we asked if the formation of spontaneous RecA-K250N or RecA-E96D foci depends on the ssDNA assembly mediators RecBCD and RecFOR. In contrast to damage-induced RecA-WT foci, which are fully dependent on these assembly proteins (as described above), we observed very similar frequencies of foci formed by either RecA-K250N or RecA-E96D in the *recBC sbcBC recF* strain as compared to wild type (Figure 5A, Figure 4C). These findings suggest that the spontaneous foci observed form by a pathway that is different from the normal pathway for assembly of RecA at sites of ssDNA.

We next asked if the spontaneous foci formed by the ATPase defective mutants involve ssDNA replication intermediates; most tracts of ssDNA that form in undamaged growing cells are associated with replication intermediates. We therefore asked if blocking replication initiation with a *dnaAts* mutation would alter the number of spontaneous foci observed with the ATPase-defective RecA mutants, as would be expected if the foci were forming at sites of replication. In order to determine if a particular cell was replicating or not, we double immunostained *dnaAts* spread nucleoids for SSB and RecA. We prepared nucleoids from *dnaAts* cells arrested at 42°C to detect replication-associated SSB foci and compared the results to those from cells incubated at 30°C, the permissive temperature for *dnaAts*. At 30°C, we detected an overall average of 5.2 ± 0.4 SSB foci per nucleoid (Figure 6A); 96% of nucleoids contained at least one SSB focus (Figure 6B). The *dnaAts* cells at the restrictive temperature displayed only 0.3 ± 0.1 SSB foci per nucleoid, 17-fold less than the number seen in the low temperature control. These results show that spontaneous SSB foci are associated with on-going replication and provide a new method for cytological detection of replication intermediates in *E. coli*. After cells were arrested, they were chilled on ice, harvested by centrifugation at 4°C, and re-suspended in medium containing arabinose that had been pre-warmed to 42°C, or to 30°C for the control culture. A second set of spread nucleoids was then prepared following a 30-minute incubation at the same temperatures to induce expression of RecA variants. Nucleoids prepared after the induction protocol displayed significantly more SSB foci than did the samples taken before RecA induction, suggesting the block to replication was not fully maintained during the media transfer procedure because the temperature was reduced during the procedure. This feature of the method provides a convenient control that allows us to compare cells from the same sample that maintained replication arrest with ones that did not. We divided the focus counts from the total population of nucleoids examined into two subpopulations, one in which the spread nucleoids contained no SSB foci, and one in which the nucleoids displayed at least one SSB focus. We then compared the frequency of RecA foci in the two subpopulations. We detected no significant difference in the average frequencies of RecA-K250N (P=0.76, Mann-Whitney U test); the SSB focus-positive population had 1.6 ± 0.3 foci per nucleoid, and the focus-negative population had 1.5 ± 0.3 foci per nucleoid. Similar results were obtained from RecA-E96D-expressing nucleoids prepared after incubation at 42°C; there was no significant difference in the average frequencies of RecA-E96D foci between SSB-positive and SSB-negative populations (P=0.11, Mann-Whitney U test), and the average number of RecA-E96D foci in the SSB-positive population was 2.3 ± 0.3 foci per nucleoid in SSB-positive population and 1.6 ± 0.3 foci per nucleoid in SSB-negative population. These results indicate that formation of foci by the two ATPase defective variants is independent of DNA replication. We also compared the numbers of RecA foci seen in 30°C control culture (Figure 6C). In this control, the frequency of induced foci was significantly lower than at 42°C, suggesting that focus induction may be slower at 30°C than at 37°C, the temperature used in the experiments described above.

**Figure 6.**
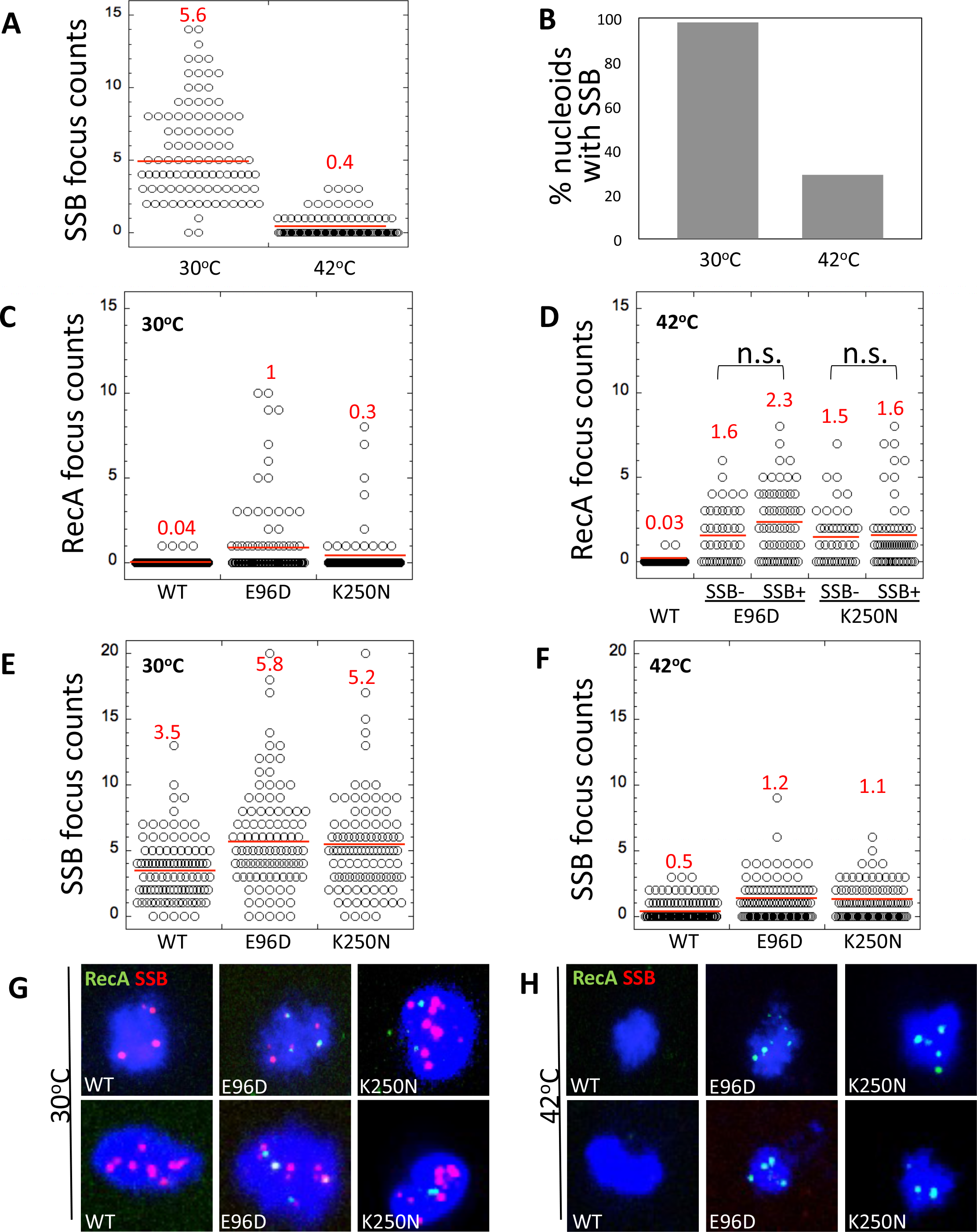
Analysis of RecA ATPase defective variants in *dnaAts* cells (DB82). A. Average number of SSB foci per nucleoid in cells prior to RecA induction. B. Percentage of nucleoids that contain SSB foci prior to RecA induction. C. RecA focus counts after RecA induction at 30°C. D. RecA focus counts after RecA induction at 42°C. E. SSB focus counts after RecA induction at 30°C. F. SSB focus counts after RecA induction at 42°C. Spread nucleoids were prepared, foci from individual nucleoids were counted, and data plotted as described in legend of Fig. 4. G. Representative nucleoids from the 30°C experiment. H. representative nuclei from the 42°C experiment. RecA (green), SSB (red); DNA was stained with DAPI (blue).

## DISCUSSION

In this study we show that RecA’s ATPase activity is required to prevent accumulation of cytologically-detectable amounts of RecA protein on *E. coli* chromosomal DNA in the absence of exogenous DNA damage. This study follows from the groundbreaking work of Campbell and Davis who identified toxic forms of RecA and biochemically characterized RecA-E96D (24, 25). Our work extends the conclusions of that previous work in two important ways. We first show that ATPase deficiency alone is sufficient to account for the toxicity observed; RecA-K250N is ATPase defective, and binds both ssDNA and dsDNA more slowly and with lower affinity than RecA-WT, but is nearly as toxic as RecA-E96D, which displays faster than normal DNA binding.

We also provide key new information regarding the cause of toxicity conferred by ATPase defective proteins. Campbell and Davis showed that strains expressing RecAE96D were capable of promoting homologous recombination, leading them to favor a model in which the mutant protein’s toxicity is caused by the failure of the mutant protein to dissociate from recombination intermediates (24). Although our D-loop experiments provide additional evidence that RecA-E96D is proficient in recombination, and also suggest the same is true for RecA-K250N, the demonstration of recombination proficiency does not imply that a defect in the recombination pathway is responsible for toxicity. This issue was recognized by Campbell and Davis; they acknowledged the possibility that dsDNA binding might be responsible for toxicity (25). The experiments described here implicate direct, non-recombinogenic, dsDNA binding as the cause of toxicity.

Our method of immunostaining spread nucleoids allows us to visualize complexes of nucleoid-bound RecA. The method allowed us to show that cells expressing DNA binding proficient, ATP hydrolysis-defective forms of RecA accumulate spontaneous complexes that are not observed in cells expressing the wild type protein. In contrast to the damage-induced structures formed by RecA-WT, the spontaneous structures formed by the toxic mutants do not depend on recombination proteins that are normally required for loading RecA on tracts of ssDNA. Furthermore, the foci do not depend on ongoing replication, which is the predominant process responsible for formation of spontaneous ssDNA tracts in the cell, as vividly illustrated by our nucleoid spreading method combined with staining for SSB. Consistent with our *in vivo* observations, we provide biochemical evidence that the dsDNA binding activity of pure RecA-WT is more prominent than revealed by earlier studies that employed different methods of detection (13, 49); using fluorescence polarization to detect bound DNA, we observed that RecA-WT bound dsDNA with nanomolar affinity (*K_d_* ~ 400nM). Although it is unknown what fraction of the highly-compacted chromosomal DNA is accessible for RecA binding in living *E. coli* cells, it is worth noting that the cellular concentration of RecA (16 µM assuming 10^4^ molecules/cell and a cell volume of 1 fL) exceeds the *K_d_* for dsDNA binding reported here. This implies that RecA will bind available tracts of dsDNA in the cell. Taken together, our observations make it extremely unlikely that the nucleoid bound structures seen with RecA-E96D or RecA-K250N reflect a ssDNA bound form.

We also note that the viability of *recA* null mutants also makes it very unlikely that the 3-4 logs of kill seen upon induction of RecA-K250N or RecA-E96D relates to defective recombination or DNA repair. A recent estimate of the fraction of cells that suffer unrepaired DSBs in *recB* mutants is 0.18 (50). Therefore, if the profound toxicity associated with RecA-E96D and RecA-K250N were a consequence of blocking repair of a sufficient number of lesions to account for 3 logs of kill, an alternative repair mechanism must normally function to repair those lesions in a *recA* null mutant. However, the results of a search for mutations that confer synthetic lethality with a *recA* null mutation argue against this possibility (51). The mutations identified involved pathways that prevent spontaneous DNA damage, not pathways that repair damage. These considerations contribute to our argument that the toxicity caused by ATPase-defective forms of RecA does not reflect a block to the normal, RecA-dependent DNA recombinational repair pathway, but instead reflects off-pathway activity.

Our observations regarding the function of RecA’s intrinsic ATPase provide an interesting comparison to RecA’s structural and functional homologs in eukaryotes, Rad51 and Dmc1. Two very different ATP hydrolysis-dependent mechanisms appear to serve the common function of dissociating off-pathway complexes formed by this family of strand exchange proteins (Figure 7 B, C). Both Rad51 and Dmc1 bind dsDNA efficiently *in vitro* and both form off-pathway complexes on dsDNA *in vivo* (26-28). Unlike the intrinsic ATPase activity of RecA, the corresponding activities of Rad51 and Dmc1 are not sufficient to prevent accumulation of toxic complexes; Rad54-related translocases are required to disassemble those complexes via ATP hydrolysis-driven translocation along dsDNA (29, 30). The fact that the intrinsic ATPase activity of Rad51 and Dmc1 is about 30 to 60 fold lower than that of RecA (*k*_cat_ of 0.5-1.0 min^−1^ vs. 30 min^−1^) (31, 52) could reflect evolution of a dissociation mechanism dependent on intrinsic ATPase activity to one dependent on translocases.

**Figure 7.**
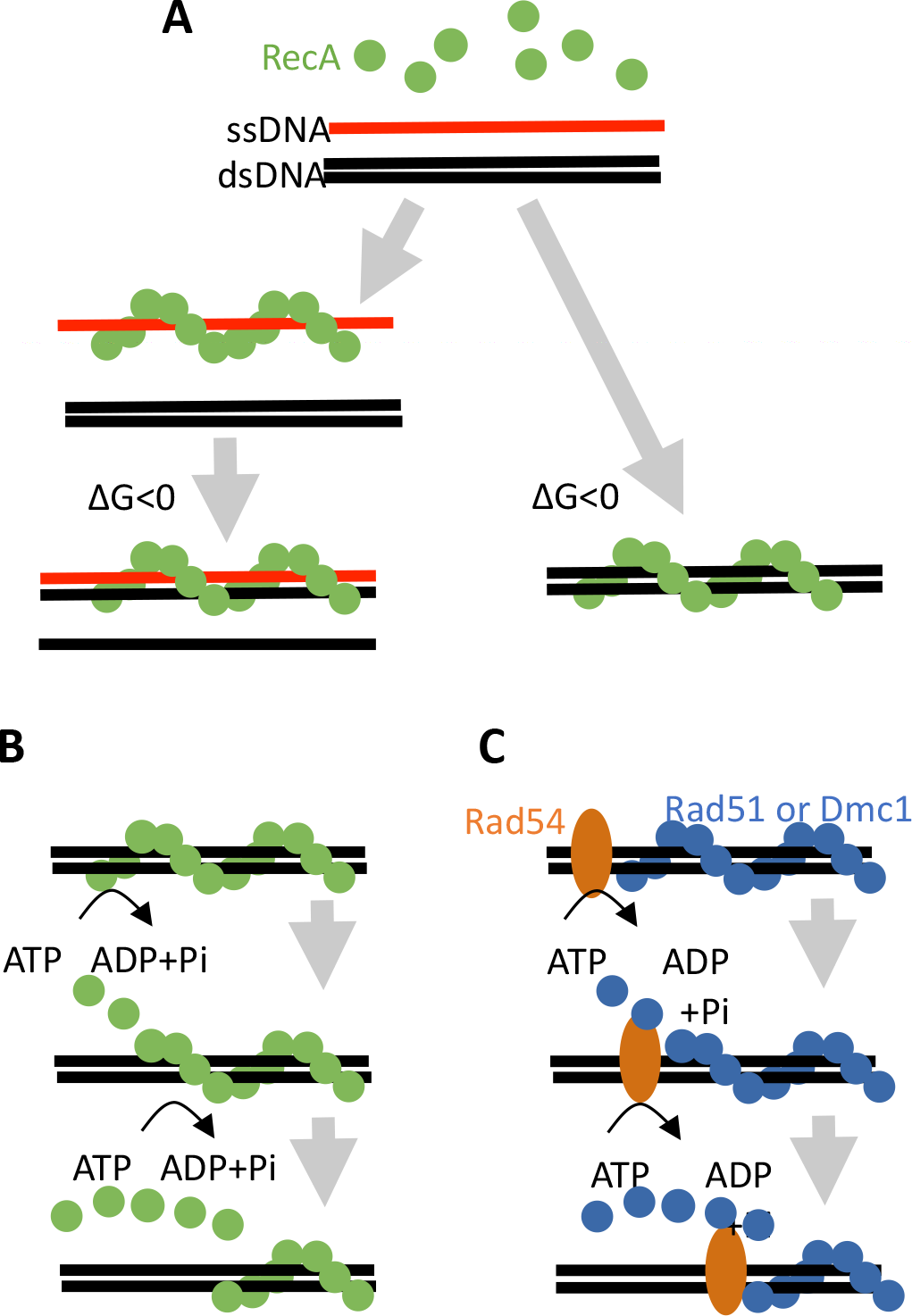
Model for the *in vivo* function of the ATP consumption associated with RecA family strand exchange protein activity. A. RecA can bind either ssDNA or dsDNA. If it is recruited to ssDNA, if forms a helical filament by binding to Site I (left side). The helical filament searches for and forms a synaptic complex with a homologous region of dsDNA. Strand exchange then occurs with the original ssDNA remaining in Site I. The strand exchange process is driven by product stability, <u>not</u> ATP hydrolysis. RecA Site I can also bind undamaged dsDNA segments of the genome (right side). This binding activity is a consequence of the mechanism of strand exchange which forms hybrid dsDNA bound to Site I. B. In *E. coli*, the intrinsic ATPase activity converts the high affinity binding form, RecA-ATP, to the low affinity binding form, RecA-ADP driving dissociation from either the hybrid dsDNA product of strand exchange or from undamaged dsDNA. C. In eukaryotic cells, dissociation of Rad51 or Dmc1 from dsDNA requires the action of a RAD54 family dsDNA-specific translocase which uses the energy of ATP hydrolysis to dissociate the protein from strand exchange products or regions of undamaged dsDNA.

If RecA’s direct dsDNA binding activity results in formation of toxic complexes, why have cells not evolved to eliminate this activity? As previously proposed for the eukaryotic homologs (28), we propose that the off-pathway activity of RecA is an unavoidable consequence of the reaction mechanism through which RecA promotes strand exchange; strand exchange is driven by product stability in a mechanism that involves the conversion of a ssDNA-ATP-RecA filament, in which the DNA occupies the high affinity binding site (Site I) of each protomer, to a dsDNA-ATP-RecA filament in which the heteroduplex dsDNA product is also bound at Site I, i.e. the original ssDNA strand acquires a complementary partner while remaining bound at Site I (Figure 7A). For this mechanism to function, RecA filaments must be capable of binding dsDNA at Site I and that activity has the consequence that the protein binds dsDNA directly. Previous work indicated the hydrolysis of RecA-ATP-dsDNA to RecA-ADP-dsDNA resulted in dissociation of RecA from heteroduplex DNA following strand exchange (17). The results presented here are consistent with the idea that ATP hydrolysis promotes dissociation from hybrid DNA recombination intermediates, but we argue that a fully analogous and essential mechanism acts to prevent accumulation of RecA-ATP-dsDNA complexes that form by direct assembly of RecA on dsDNA. In this view, ATP hydrolysis serves an ATP hydrolysis-dependent error-correction function that allows cells to discard the dead-end and potentially toxic complexes that result from direct binding to dsDNA. This type of error-correction process differs from the classically-defined kinetic proof-reading mechanism in two ways (53). First, the “incorrect” substrate, a tract of protein-free dsDNA, is likely to be in vast excess to the correct substrate, ssDNA, in the cell. Second, the “incorrect” substrate is structurally identical to the reaction product, a RecA-ATP-dsDNA filament. The strand exchange mechanism represents a constraint on evolution of the protein, hence the energetic cost of error correction is required. We argue the fitness benefit derived from homology-driven DNA repair outweighs the expense of error correction.

Here we have provided evidence for an *in vivo* function of the ATPase activity of RecA. We note that a popular hypothesis maintains that the ATPase activity of RecA also serves to allow its dissociation from products of recombination (12, 16). Although we believe this hypothesis is likely to be correct, we emphasize that, to our knowledge, it has yet to be tested by *in vivo* experiments. The need for such experiments is highlighted by the fact that RecA’s preferential binding to ssDNA does not account for its recruitment to tracts of ssDNA *in vivo*; protein-protein interactions involving SSB and mediator proteins are required (7). Therefore, the hypothesis that dissociation of RecA from recombination products is governed by its ATPase activity could be incorrect. The same is true in the eukaryotic systems where a role of translocases in dissociating Rad51 and Dmc1 from recombination products has yet to be demonstrated. In this context it should be noted that there is a competing hypothesis for Dmc1. The proteasome localizes to sites of recombination on meiotic chromosomes, raising the possibility that Dmc1 is removed from recombination products by programmed degradation (54). *In vivo* experimentation is required to determine how strand exchange proteins are removed from recombination products in cells.

Our effort to study the dynamic interaction of RecA with DNA led us to develop a simple nucleoid spreading/immunostaining method to detect and quantitate complexes of RecA associated with nucleoids. This method complements the powerful live cell methods that have been described and offers advantages those systems do not currently provide (50, 55, 56); 1. it provides much higher spatial resolution that enhances detection of closely-spaced, but not molecularly bound complexes. 2. it does not require tagging of protein with fluorescent domains that often compromise protein function, especially in the case of strand exchange proteins; 3. it removes background signal contributed by protomers not bound to the nucleoid, improving detection of complexes that contain small numbers of molecules; and 4. it can be used with standard wide-field or confocal microscopes with straightforward and fast sample preparation. As is the case for other widely used chromosome spreading methods, one has to be aware that a significant fraction of DNA-bound protein may be lost during the spreading procedure. The same is true for protein-protein contacts between adjacent complexes that bind more tightly to DNA than to each other. Nonetheless, the method allows one to detect changes in the number and staining intensity of nucleoid bound structures under various growth conditions and genetic backgrounds. It should be noted that the utility of nucleoid spreading is not limited to the study recombination proteins. Here we show it is possible to detect SSB containing replication intermediates and it may also be possible to detect proteins engaged in additional pathways of DNA metabolism. A further advantage of nucleoid spreading is that the samples that are particularly amenable to analysis via super-resolution light microscopy. Here we used STED microscopy to study the size of damage-induced RecA complexes. When taken together with known RecA filament structural and flexibility properties, our results suggest that both the DNA damage-dependent and the spontaneous RecA structures we observe are rarely as long as 400 and often equal to or shorter than 100 nt (or bp). This finding is significant, since little prior information is available on the lengths of physiologically-active RecA filaments, despite a great deal of biochemical data on the properties of RecA filaments formed on a wide length range of DNA substrate lengths (47, 57, 58). We note, however, that our estimate of filament length assumes that the persistence length of RecA filaments *in vivo* corresponds to that *in vitro*, if filaments are folded by some as yet to be discovered mechanism, they could be longer than our current estimate suggests.

## ACKNOWLEDGEMENTS

We thank Susan T. Lovett, Ron Davis, and David Bates for strains and plasmids. We are grateful to Ute Curth for the gift of SSB antibodies. We thank Benjamin P. Weissman for assistance in analysis of biochemical data. We also thank Alex Ruthenburg for critical reading of the manuscript.

